# An *in silico* approach to identification, categorization and prediction of nucleic acid binding proteins

**DOI:** 10.1101/2020.05.05.078741

**Authors:** Lei Xu, Shanshan Jiang, Quan Zou

**Affiliations:** School of Electronic and Communication Engineering, Shenzhen Polytechnic, Shenzhen 518000, China; School of Software & Microelectronics, Peking University, Beijing 102600, China; Institute of Fundamental and Frontier Sciences, University of Electronic Science and Technology of China, Chengdu 651004, China

## Abstract

The interaction between proteins and nucleic acid plays an important role in many processes, such as transcription, translation and DNA repair. The mechanisms of related biological events can be understood by exploring the function of proteins in these interactions. The number of known protein sequences has increased rapidly in recent years, but the databases for describing the structure and function of protein have unfortunately grown quite slowly. Thus, improving such databases is meaningful for predicting protein-nucleic acid interactions. Furthermore, the mechanism of related biological events, such as viral infection or designing novel drug targets, can be further understood by understanding the function of proteins in these interactions. The information for each sequence, including its function and interaction sites, were collected and identified, and a database called PNIDB was built. The proteins in PNIDB were grouped into 27 classes, such as transcription, immune system, and structural protein, etc. The function of each protein was then predicted using a machine learning method. Using our method, the predictor was trained on labeled sequences, and then the function of a protein was predicted based on the trained classifier. The prediction accuracy achieved a score of 77.43% by 10-fold cross validation.

**Availability and Implementation:** PNIDB is now fully working and can be freely accessed at: http://server.malab.cn/PNIDB/index.html. All the data are publicly available for non-commercial use, distribution, and reproduction in any medium.

## 1 Introduction

As the chief actors within the cells, proteins are involved in many essential activities in the cell, and the interactions between proteins with nucleic acid are extremely important for many cellular processes, such as transcription, translation, and DNA repair. Therefore, the study of protein-nucleic acid binding activities can help with understanding protein interaction networks or even the mechanism of related cellular processes. With the development of sequencing technology, the amount of protein sequence information has increased rapidly over recent years, but the growth rate of databases describing the structure and function of proteins has been very slow, and cannot match the growth rate of protein sequence databases. Thus, it is essential to narrow the gap between sequence database size and functional database size by characterizing nucleic acid binding proteins and their functional groups. Therefore, a database called PNIDB (Protein-Nucleic acid Interactions Database, PNIDB for short), in which DNA-binding proteins and RNA-binding proteins are denoted by gene ontology, was built. Sequence information was extracted, and an efficient classification method for predicting DNA-binding and RNA-binding proteins is proposed in this article. The process of our work is summarized in Fig. 1.

**Fig. 1.**
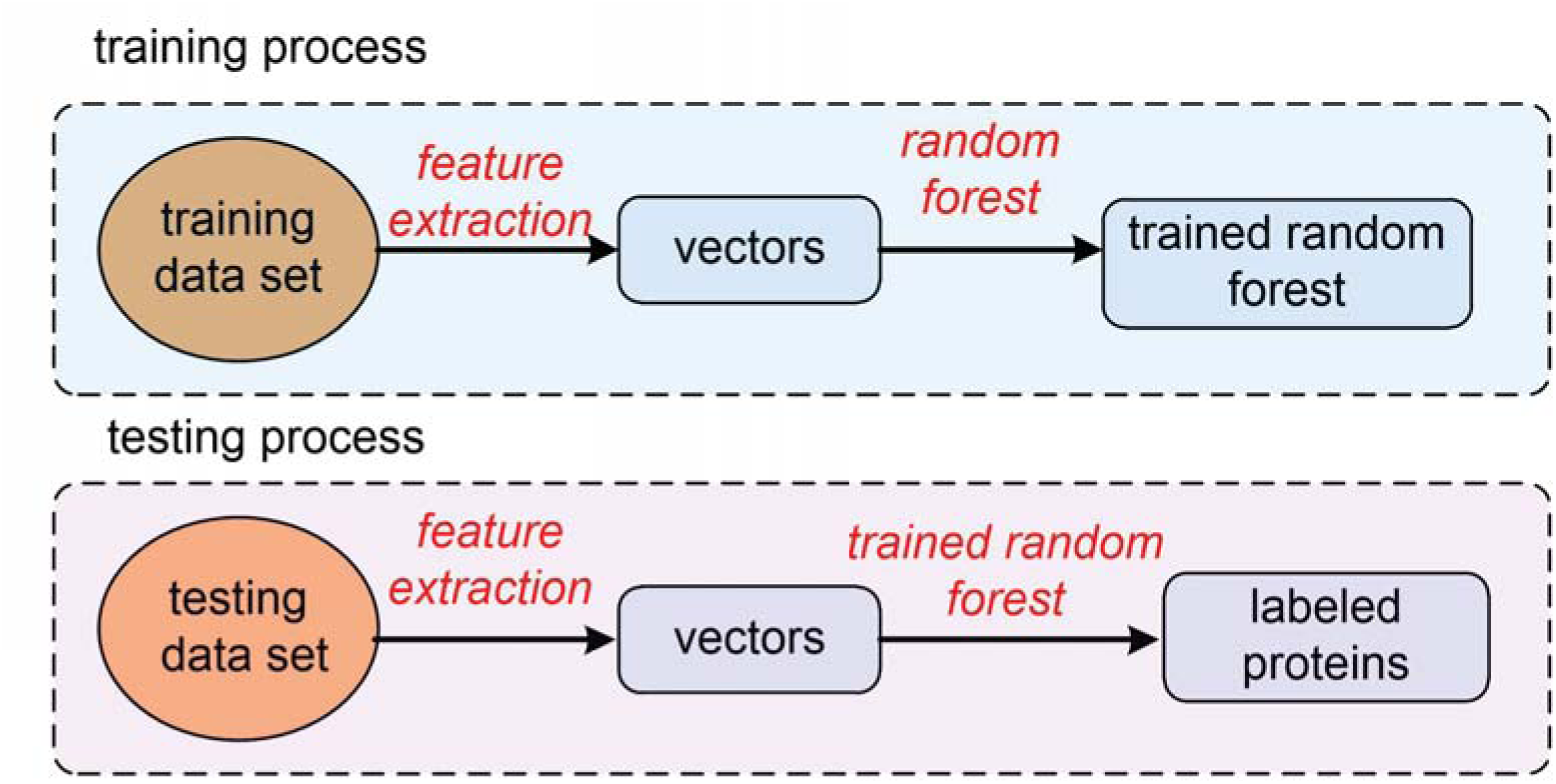
The process of protein prediction.

As shown in Fig. 1, the data set was built including DNA-binding and RNA-binding proteins. Each sample was represented by a 473-dimensional vector describing the sequence information and secondary structural information. The parameters of a random forest model were trained using the training samples. The testing samples were then classified by the random forest model with the previously learned parameters.

There are studies focusing on identifying nucleic acid binding proteins and non-nucleic acid binding proteins, and the accuracy of existing methods has been improved over time (Qu & Zou, 2018). However, these methods cannot distinguish the type of binding proteins, such as DNA binding proteins or RNA binding proteins, so it is meaningful to be able to predict the functions of these proteins. Based on existing research, our work focused on identifying and distinguishing DNA-binding proteins and RNA-binding proteins. Since the secondary structure of RNA is diverse, it is difficult to identify common characteristics of RNA binding proteins. As a result, the problem of identifying RNA binding proteins has been rarely considered in previous studies. Thus, using current methods, RNA binding proteins are likely to be recognized as DNA binding proteins. In fact, the function of DNA binding proteins is quite different from the function of RNA binding proteins; thus, the study of distinguishing DNA binding proteins and RNA binding proteins should be considered. In our current study, RNA binding proteins were divided into different classes depending on their characteristics, and the identification of DNA binding proteins was discussed. Furthermore, for the purpose of revealing the biological functions of binding proteins in cellular activities, a protein-nucleic acid binding database called PNIDB was created and can be accessed online. The information, such as functional classification of protein chain and binding events at the sequence level, are all described in PNIDB. Moreover, the database can predict protein-nucleic acid binding events with the information given by PNIDB.

In contrast to previous work, the nucleic-acid binding proteins were further divided into RNA binding proteins and DNA binding proteins in the proposed method, and both of them were identified by gene ontology in PNIDB. The rationale was that the functions of proteins can be learned thoroughly when the proteins are identified precisely. The contributions of our work include:

1. A database describing protein-nucleic acid interactions, known as PNIDB, which can be accessed by researchers at the website http://server.malab.cn/PNIDB/index.html. The information provided by PNIDB can help reveal the biological functions of binding proteins in cellular activities. The proteins in PNIDB are labeled by gene ontology identifiers.
2. An efficient classifier for predicting DNA binding proteins and RNA binding proteins was proposed based on sequence information and secondary structural information, and the accuracy of the proposed method achieved a correlation of 77.43%, which outperforms other methods. The experimental results demonstrated that the combined information could improve the prediction accuracy of our method.
3. A web server for the prediction of protein-nucleic acid was also developed. The web server was used to predict the function of proteins, which can help researchers studying protein-nucleic acid interactions.

In the rest of the paper, the usage manual PNIDB is introduced in Section 2. Section 3 introduces the methodology for the classification of functional proteins in detail. The experimental results are reported in Section 4. Finally, conclusion are made in Section 5.

## 2 Usage of PNIDB

There are other databases that provide information regarding protein-DNA/RNA interactions. For example, the Protein-DNA Interface database (PDIdb) (Norambuena, 2010) is a repository containing structural information for 922 protein-DNA complexes with a resolution of 2.5 Å or more (while in fact there are 2396 this kind of complexes in the databaset). The Nucleic acid-Protein Interaction database (NPIDB) (Kirsanov, Zanegina, Aksianov, Spirin, & Alexeevski, 2012) (Olga et al., 2015) contains structural classifications and detailed information on both DNA-protein and RNA-protein complexes extracted from PDB. The current version of NPIDB contains 5046 structures, while PNIDB contains 6228 PDB structures overall. In contrast to the above databases, proteins are denoted by gene ontology in PNIDB.

PNIDB provides detailed atom-based interaction information. The significance of PNIDB is that it specifically focuses on sequence-level annotations and provides functional clustering, which should be of benefit for sequence-based research and functional prediction of protein-nucleic acid interactions.

PNIDB is a repository of protein-nucleic acid interaction information derived from 6,798 nucleic acid containing structures collected from the Protein Data Bank (Burley et al., 2015). The Bio3d (Skjaerven, Yao, Scarabelli, & Grant, 2014) package was used to read, analyze and manipulate PDB structures in R (http://www.R-project.org/). For each PDB file, we identified the proteins, and the nucleic acid chains were then extracted. Moreover, possible binding residues of the protein chain, which were defined as the residues with at least one atom within 5 Å from any nucleic acid atom, and corresponding binding nucleotides of the nucleic acid chain were also calculated in PNIDB.

Protein chains in interaction pairs were classified according to their mmCIF keywords, interaction type and Gene Ontology (Ashburner, Ball, Blake, Botstein, & Cherry, 2000) terms. There were a total of 84,753 chains extracted from those structures, in which 20,927 chains contained nucleic acid binding residues. All the protein chains were clustered into 27 functional groups, with 17 kinds of DNA binding proteins and 10 kinds of RNA binding proteins. Moreover, each protein chain in these interactions was linked to their respective accession numbers from UniProt as well as the corresponding InterPro identifiers (Finn, 2017) and GO identifiers (Ashburner et al., 2000; Carbon, 2017) mapped from the SIFTS project (Velankar, 2013).

For convenience, the residues and nucleotides are cited by their relative position in the sequence of their separate chains. In addition, the 2D and 3D visualization interfaces are provided online. Fig. 2A and Fig. 2B show the interfaces of the 2D and 3D visualization, respectively, in PNIDB.. In Fig. 2B and 2D, the visualization interface focuses on nucleic acid sequences. The binding protein residue and the position is highlighted in the figure. The residues exceeding 3.9 Å are considered binding residues.

**Fig. 2.**
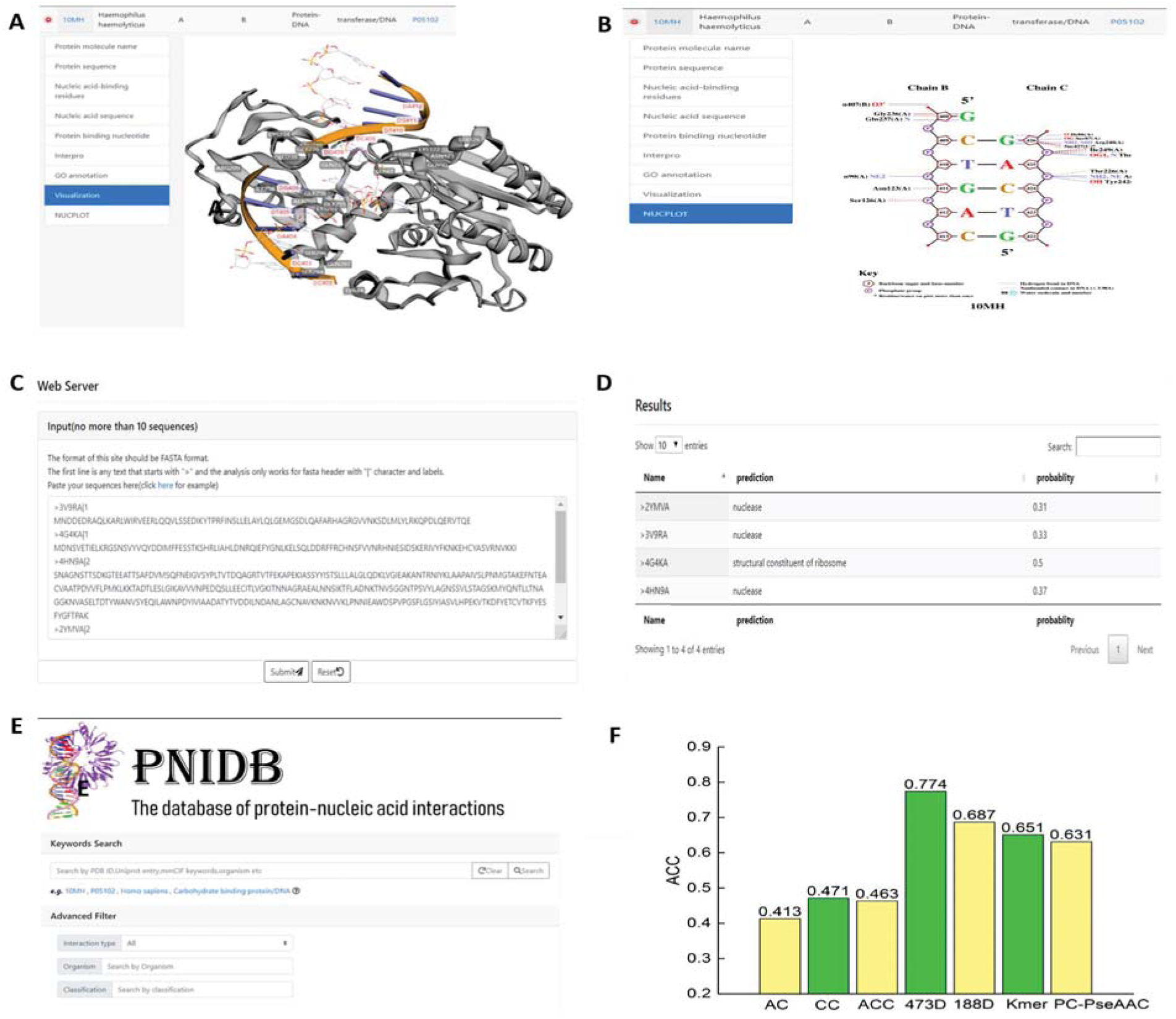
(A) 3D visualization interface of the interaction between a protein and a nucleic acid residue. (B) 2D visualization interface of an interaction (solid line for hydrogen bond). (C) The web server for binding protein prediction. (D) Results provided by the web-based predictor. (E) Search page on the web server. (F) Comparison with other methods by ACC. (Nucleic acids are labeled in red, while protein residues are in grey, and in the 2D visualization, the dash line denotes residues within 3.9 Å.)

A search page is also provided online, and users can search by keywords. A quick search and advanced search were implemented. In quick search mode, users can start a search by specifying a keyword, such as a PDB ID (Burley et al., 2015) organism, interaction type, classification or Uniprot accession number (Rolf et al., 2004). In the advanced search mode, for the convenience of other researchers, a web server for binding protein prediction was developed. The web server can handle up to 10 fasta sequences at the same time. Then, the results are returned by email. The web server is shown in Fig. 2C. The results predicted by a learned classifier are shown in Fig. 2D.

The search page of PNIDB is provided, which is shown in Fig. 2E. There are three options, “search by interaction type”, “search by organism” and “search by classification”. Users can search proteins by describing the requirements. Moreover, proteins can be searched by combining several parameters simultaneously. The matching results will be retrieved when the requirements are submitted. Furthermore, more information can be referred using PNIDB, such as the molecule name of the protein chain, the sequence of the protein chain, the binding residues of the protein chain, the sequence of the nucleic acid chain, the binding nucleotides of the nucleic acid chain, the corresponding InterPro IDs (Finn, 2017) of the protein chain, the GO identifiers of the protein chain, and the 3D visualization with labels of contacting residues and nucleotides based on 3Dmol.js (Rego & Koes, 2015). The schematic diagrams of protein-nucleic acid interactions based on NUCPLOT (Luscombe & N., 1997) can be obtained by clicking the tab on the webpage.

For the convenience of related study, all binding residues/nucleotides were renumbered according to their corresponding chain sequence. In addition, users can also browse the interactions in the browse page by selecting specific classifications of the protein chains in the menu on left side.

## 3. Methodology

The method for predicting the function of proteins is described in this section. First, the benchmark dataset used is introduced in Section 3.1. Then, the method of feature extraction is described in Section 3.2. Lastly, the classification of the binding proteins is illustrated in Section 3.3.

### 3.1 Benchmark Data Set

The data used in this work was selected from the SwissProt data set (https://web.expasy.org/docs/swiss-prot_guideline.html). The data in SwissProt contained GO protein sequences, which were selected from the UniProt data set (https://www.uniprot.org/) with high confidence. The SwissProt dataset was composed of DNA binding proteins and RNA binding proteins. The DNA binding proteins had non-IEA source gene ontologies. The sequences that were more similar were removed using CD-HIT(Li & Godzik, 2006). The similarity degree between the sequences used in our experiments was less than 30%. The benchmark dataset contained five classes. The benchmark dataset used in these experiments is summarized in Table 1.

**Table 1.** Sample detail of benchmark dataset

| Class | No. of samples |
| --- | --- |
| DNA binding transcription factor | 200 |
| Helicase activity | 256 |
| Nuclease | 200 |
| RBP spliceosomal complex | 199 |
| RBP structural constituent of ribosome | 213 |

### 3.2 Feature Extraction

In the literature, residue sequences are usually represented by a vector ***v*** before the process of prediction. An efficient feature set is expected to distinguish positive samples and negative samples with high accuracy. The quality of a feature set is critical to the performance of any predictor. In our method, the sequential evolution information, as well as the local and global secondary structural information, were combined for representing a protein sequence ***S***. The features used in our method were extracted from sequence ***S***. The features include PSI-BLAST features (Altschul et al., 1997) and PSI-PRED features (Jones, 1999). PSI-BLAST describes the evolutional information, and the secondary structural information is shown by PSI-PRED. The combination of these features has been previously used for protein fold prediction (Wei, Liao, Gao, & Zou, 2015).

A protein sequence *S*_*L*_ is denoted as *S*_*L*_={R_1_,R_2_,…,R_L_}, and *L* is the length of the residue. PSI-BLAST is based on the protein database *nrdb90* (L. Holm, 1998) and a position-specific score matrix (PSSM). A PSSM is a matrix with L× 20, written as Eq. (1):

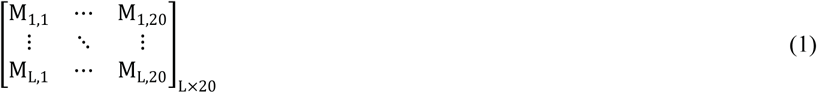

M_i,j_ is denoted as the score of the residue at the *i*th position of *S*_*L*_ being mutated to the residue type *j* during an evolutionary process. The 20 features are computed by the average value of each column (Eq.(2)):

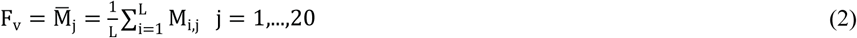

The evolution information is also used for describing a sequence. During the evolutionary process, an amino acid located at the *i*th position in the residue may be mutated to type *j*, and the score of it is denoted as ***M***_***ij***_ (Eq. (3)):

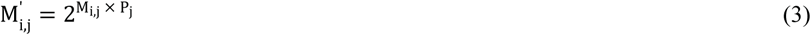

***P***_***j***_ represents the background frequency of residue type *j*, and the background depends on the average occurrence frequency of all 20 amino acid in each sequence of the protein database *PDB25* (Sussman et al., 2010).

δ_i_ is represented as the residue located at the *i*th position in an original sequence, ***S***_***L***_, replaced by the δ_i_th amino acid in the amino acid alphabet. The sequence ***S***_***L***_ is transferred into a consensus sequence ***S***_***con***_ by using δ_i_. The frequency of each δ_i_ in the sequence is denoted as a feature. Thus, there are 20 amino acids, so 20 features are extracted from each sequence.

***R***_***i***_ represents the *i*th residue of a peptide. There are 20 types of native amino acids, which means that there are 20 possible values for R_i_. Thus, the number of two consecutive amino acids R_i_R_j_ is 400 (20 × 20), meaning that the number of dimensions describing the occurrence frequency of two constitute amino acids is 400. The 400-dimensional features have been widely used in the literature of bioinformatics, such as Alzheimer’s disease identification (Xu, Liang, Liao, Chen, & Chang, 2018) and detection of anticancer peptides (Xu, Liang, Wang, & Liao, 2018). Thus, sequence information can be revealed using these 400-dimensional features. In contrast to using 400-dimensional features, the features used in our work were based on the consensus sequence S_con_. The frequency of δ_i_δ_j_ is denoted as a feature. The frequency of the occurrence of δ_i_δ_j_ is denoted as a feature in ***v***. Therefore, another 400 features are extracted from each sequence.

PSI-PRED-based features have been widely used in secondary structure prediction. The features include six structure-sequence based features and ***a***×3^n^+3^n^ structure probability matrix-based features. The value of ***a*** is set to be 8, and ***n*** is 1. Thus, there are 33 PSI-PRED features. Therefore, there are 473 (20+20+400+33) features used to represent a sequence ***S***_***L***_ in total.

### 3.3 Classification

Support vector machine, naive Bayes and ensemble methods have all been widely used in bioinformatics, such as prediction of tumor detection (Tang, Wan, Yang, Teschendorff, & Zou, 2018), function prediction of proteins, and disease detection (Xu, Liang, Liao, Chen, & Chang, 2019). The performance of a predictor is also related to the classifier used. Thus, an efficient classifier is critical for the performance of a computational predictor. In our work, a random forest model was used to predict the function of proteins.

Random forest models are a type of ensemble classifier. The key idea of random forest model is that a number of decision trees are used together for prediction. The decision trees are trained by the datasets, which are built based on bagging. Each decision tree makes a decision, and the final decision is made by a voting process. A sample is then classified into the class with the most votes. The process of a random forest model is shown in Fig.3.

**Fig. 3.**
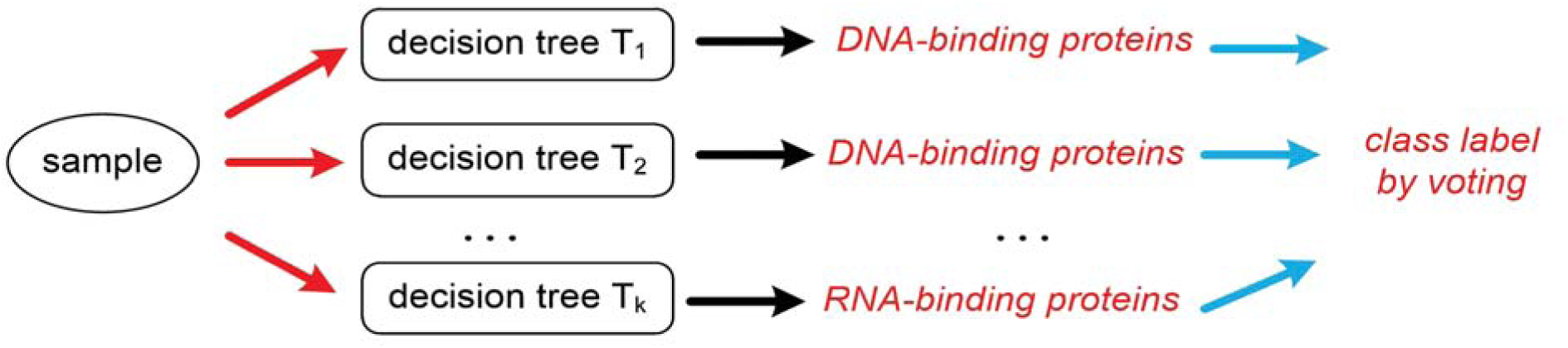
Process of random forest model generation.

## 4 Results

To demonstrate the efficiency of our proposed model, our proposed method is compared with other methods, which have been used widely in the literature.

188D is proposed to extract features from a sequence (Cai, Han, Ji, Chen, & Chen, 2003). The amino acid composition, distribution and physicochemical property are described in 188D. This method has been used in bioinformatics, such as for the identification of antioxidant proteins (Xu, Liang, Shi, & Liao, 2018).

The Kmer (k = 2) method extracts features representing the occurrence frequency of k consecutive amino acid in a residue. In our experiments, k was set to 2. The number of dimensions of Kmer features is (n-k)+1, where n is the length of residue.

PC-PseAAC was proposed based on pseudo amino acid components (PseAAC) and has been used in protein identification (Chou, 2011). The information of location residue and global residue are mixed into PseAAC in PC-PseAAC.

An autocorrelation (AC) is the correlation between any two residues with distance lag on the same properties (Liu, Liu, Fang, Wang, & Chou, 2015).

The experiments were based on a 10-fold cross validation. In a 10-fold cross validation, a dataset is divided into 10 parts. Ninety percent of samples are then used for training parameters, and the remaining 10% are considered as testing samples.

The evaluation metric was measured by accuracy (ACC), which was denoted as the rate of correctly classified samples using method ***G***. ACC has been widely used in the literature (Chen, Peng, Han, Cai, & Cai, 2018). The data set was composed of both positive and negative samples. The result set was divided into four parts, which were true positive (TP), false positive (FP), true negative (TN) and false negative (FN). TP is the number of positive samples classified correctly. FP is the number of negative samples labeled as positive samples. TN denotes the number of negative samples that were labeled correctly. FN is the number of positive samples that were recognized as negative samples. The accuracy (ACC) of method ***G*** was computed using Eq. (4):

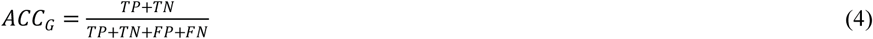

The comparison of the feature sets is shown in Fig. 2F. The experimental results show that our proposed method performed better than other feature sets. The combination of sequence information features improved the prediction accuracy. In fact, the features used in Kmer (k=2) were a part of the features in 473D. In the experimental results, the accuracy was improved by nearly 19% (((77.43–65.07)/65.07)%) when the sequences were represented by the PSI-BLAST and PSI-PRED features compared with only using the Kmer (k=2) features. The sequence information was also extracted in 188D, so the accuracy was 0.69, which outperforms other existing methods except 473D. The accuracy of AC and CC were 0.41 and 0.47, respectively. The accuracy of the combination of AC and CC was 0.4625, which was not as good as other methods, such as 188D. In the experiments, the features of AC and CC were not suitable for predicting the function of proteins. The sequence information and secondary structural information were helpful for improving the accuracy and have been used in our proposed method.

## 5 Conclusions

Due to the rapid growth in the number of protein sequences without identification of their functions, a database describing the protein-nucleic acid interactions (PNIDB) was provided in our work. The functions of sequences were labeled using GO identifiers in PNIDB. PNIDB provides a convenient and user-friendly interface to query and browse detailed information on protein-nucleic acid interactions. Different from existing databases, PNIDB focuses on both protein-DNA and protein-RNA interactions, and the functional classifications are considered at the sequence level. Moreover, a benchmark database is available for the prediction of protein-nucleic acid binding events at either the protein residue level or nucleotide level. PNIDB will also aid in the functional prediction of nucleic-binding proteins based on protein sequence, and may help for providing putative drug targets and novel therapy options. The problem of classification of DNA-binding proteins and RNA-binding proteins was also considered in this work. The sequences are represented by PSI-BLAST features and PSI-PRED features, and a random forest model was used to predict the type of protein, such as DNA-binding proteins and RNA-binding proteins. The accuracy of our proposed method was 0.774, which performs better than other methods. A web server for protein prediction was provided online for the convenience of other researchers. Above all, PNIDB labeled by gene ontology identifiers was built for describing the function of proteins, and a computational predictor was developed for classifying DNA-binding proteins and RNA-binding proteins.

## Funding

This work was supported by the Natural Science Foundation of China (Nos. 61902259, 61771331), the Natural Science Foundation of Guangdong province (grant no. 2018A0303130084), and the Science and Technology Innovation Commission of Shenzhen (grant nos. JCYJ20170818100431895).

## Conflict of Interest

None declared.

